# ADAR2-mediated Q/R editing of GluK2 regulates kainate receptor upscaling in response to suppression of synaptic activity

**DOI:** 10.1101/443010

**Authors:** Sonam Gurung, Ashley J. Evans, Kevin A. Wilkinson, Jeremy M. Henley

## Abstract

Kainate receptors (KARs) regulate neuronal excitability and network function. Most KARs contain the subunit GluK2 and the properties of these receptors are determined in part by ADAR2-mediated mRNA editing of GluK2 that changes a genomically encoded glutamine (Q) to arginine (R). Suppression of synaptic activity reduces ADAR2-dependent Q/R editing of GluK2 with a consequential increase in GluK2-containing KAR surface expression. However, the mechanism underlying this reduction in GluK2 editing has not been addressed. Here we show that induction of KAR upscaling results in proteasomal degradation of ADAR2, which reduces GluK2 Q/R editing. Because KARs incorporating unedited GluK2(Q) assemble and exit the ER more efficiently this leads to an upscaling of KAR surface expression. Consistent with this, we demonstrate that partial ADAR2 knockdown phenocopies and occludes KAR upscaling. Moreover, we show that although the AMPAR subunit GluA2 also undergoes ADAR2-dependent Q/R editing, this process does not mediate AMPAR upscaling. These data demonstrate that activity-dependent regulation of ADAR2 proteostasis and GluK2 Q/R editing are key determinants of KAR, but not AMPAR, trafficking and upscaling.

**Summary statement:** Synaptic suppression promotes proteasomal degradation of the mRNA-editing enzyme ADAR2. Decreased ADAR2 levels reduce Q/R editing of the kainate receptor subunit GluK2 leading to enhanced surface expression and homeostatic upscaling.

## Introduction

Kainate receptors (KARs) are glutamate receptors comprising tetrameric assemblies of combinations of five receptor subunits, GluK1-5, with GluK2 and GluK5 being the most abundant subunit combination (Aad et al., 2010; Kumar et al., 2011; Petralia et al., 1994). KARs can be located pre-, post- and/or extrasynaptically, where they contribute to neurotransmitter release, postsynaptic depolarisation and the regulation of neuronal and network excitability. The variety of possible subunit combinations, together with co-assembly with Neto auxiliary subunits (Griffith and Swanson, 2015), creates a wide range of possible KAR subtypes (Evans et al., 2017a).

Additional KAR diversity arises from RNA editing (Egebjerg et al., 1994; Howe, 1996) mediated by the nuclear enzyme ADAR2 that edits pre-mRNAs encoding GluK2 and GluK1, as well as the AMPAR subunit GluA2, and other non-coding RNAs (Nishikura, 2016; Sommer et al., 1991). ADAR2-mediated Q/R editing in the pore-lining region of GluK2 alters a genomically encoded glutamine residue to an arginine, changing receptor assembly efficiency, forward trafficking, calcium permeability and biophysical properties of the KARs (Egebjerg et al., 1994; Howe, 1996). More specifically, edited GluK2(R) has markedly reduced tetramerisation leading to its accumulation in the ER (Ball et al., 2010). Furthermore, GluK2(R)-containing KARs that do assemble, exit the ER, and reach the plasma membrane are calcium impermeable and have a channel conductance of less than 1% of non-edited GluK2(Q)-containing KARs (Swanson et al., 1996).

ADAR2 levels are very low during embryogenesis but increase in the first postnatal week (Behm et al., 2017) to edit ~80% of GluK2, ~40% of GluK1 and ~99% of GluA2 subunits in the mature brain (Bernard et al., 1999; Filippini et al., 2016; Paschen et al., 1997). ADAR2 knockout mice die at the early postnatal stage, but can be rescued by expressing the edited form of GluA2, demonstrating that unedited AMPARs are fatally excitotoxic (Higuchi et al., 2000). In contrast, mice specifically deficient in GluK2 Q/R editing are viable but are seizure prone and adults retain an immature form of NMDAR-independent long term potentiation (LTP) (Vissel et al., 2001). Thus, although not critical for survival, GluK2 editing plays important roles in network function and long-term potentiation (LTP). These observations have become particularly intriguing in the light of recent data from our lab showing that KARs can induce a novel form of LTP (KAR-LTP_AMPAR_) even in mature rats (Petrovic et al., 2017).

As well as directly inducing synaptic plasticity of AMPARs, KARs themselves undergo long-term depression (LTD) (Chamberlain et al., 2012) and LTP (González-González et al., 2012; Martin et al., 2008; Martin and Henley, 2004). Furthermore, it has recently been shown that KARs also undergo homeostatic plasticity (scaling) (Evans et al., 2017b) which, for AMPARs and NMDARs, is a crucial regulator of neuronal network and brain function because it constrains neuronal firing to within a tunable physiological range (Mu et al., 2003; Turrigiano, 2011; Turrigiano et al., 1998). This homeostatic control prevents runaway excitation thereby maintaining synaptic stability, critical to network formation, development and stability. Furthermore, defects in this process have been implicated in neurological diseases including epilepsy and schizophrenia (Wondolowski and Dickman, 2013).

We recently reported that, as for AMPARs, KAR upscaling can be induced by suppression of synaptic activity with TTX for 24 hours and that it is accompanied by a decrease in GluK2 Q/R editing (Evans et al., 2017a). However, the molecular and cellular mechanisms that reduce Q/R editing and whether this change in editing is a direct cause of, and sufficient to mediate, KAR upscaling are not known.

Here we show that the widely used and well-established ‘classical’ protocol for inducing homeostatic plasticity of AMPARs scaling by prolonged suppression of network activity with TTX in neuronal cultures (Turrigiano, 2011; Turrigiano et al., 1998) leads to KAR upscaling by promoting the proteosomal degradation of ADAR2. Decreased levels of ADAR2 reduce GluK2 pre-mRNA Q/R editing and lead to enhanced surface expression of GluK2-containing KARs. Importantly, this upscaling mechanism is specific to KARs as TTX did not change the editing status of GluA2. Together these data demonstrate a selective role of mRNA editing by ADAR2 in homeostatic upscaling of KARs and identify alterations in ADAR2 stability as a novel mechanism for inducing plasticity.

## Results and Discussion

### Suppression of synaptic activity decreases ADAR2 levels

Our previous results demonstrated that 24 h incubation with TTX treatment leads to reduced Q/R editing of GluK2 and KAR upscaling (Evans et al., 2017a) but the mechanisms leading to this are unknown. We therefore examined levels of the enzyme ADAR2, which mediates Q/R editing of GluK2. Chronic blockade of action potentials with TTX for 24 h decreased ADAR2 levels by ~50%, with no effect on ADAR1 levels (**Fig 1A-D**). Longer periods of TTX treatment did not decrease ADAR2 levels any further (**Fig 1E,F**), suggesting that a basal level of ADAR2 is retained even under long-term suppression of synaptic activity. Importantly, TTX treatment did not alter total levels of either GluK2 or GluK5 KAR subunits (**Fig S1A-C**).

**Figure 1:**
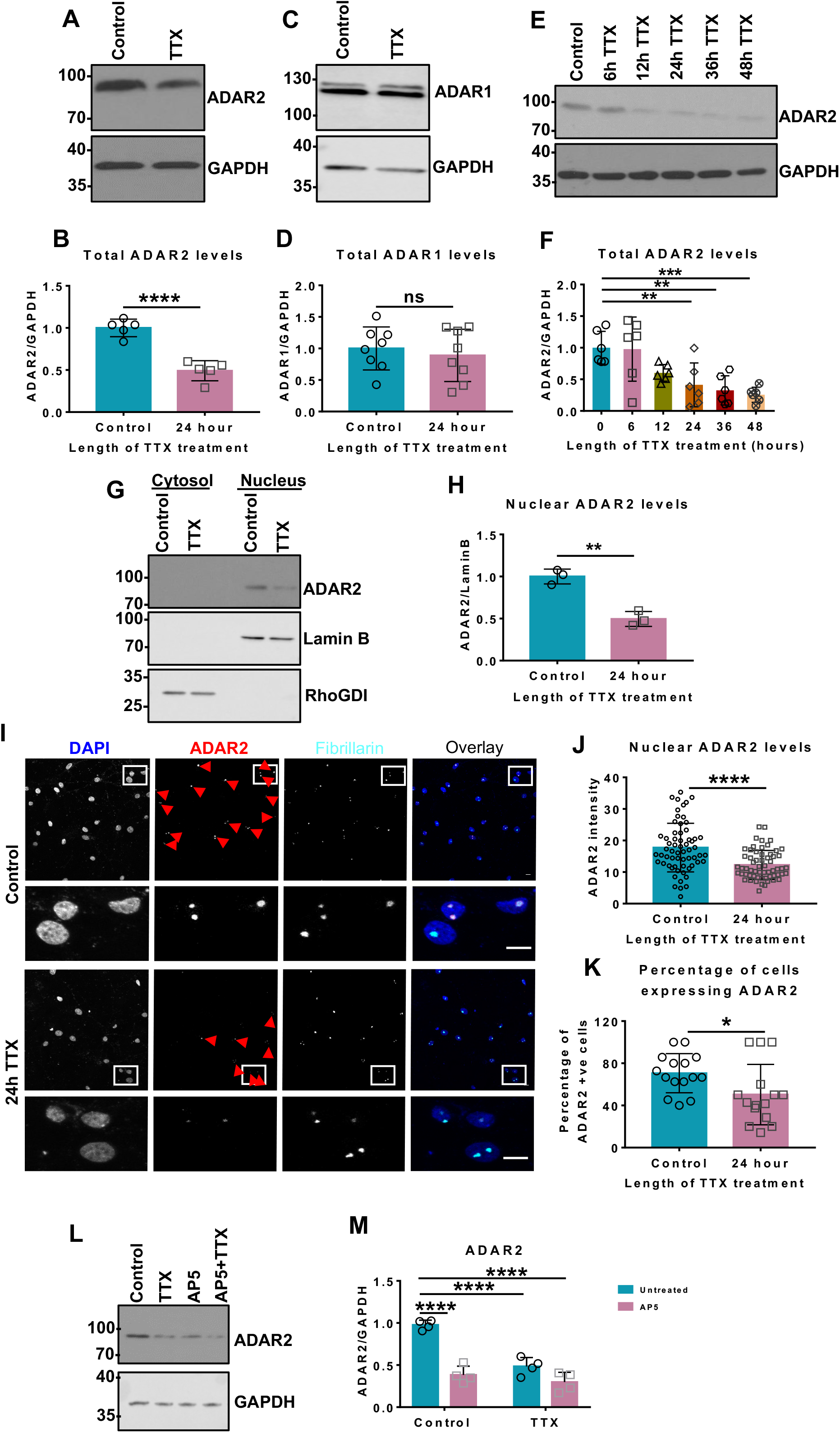
Chronic suppression of network activity decreases ADAR2 levels. A. Representative western blots of total ADAR2 and GAPDH levels in hippocampal neurons with or without 24 h TTX treatment to suppress synaptic activity. B. Quantification of (A) total ADAR2 normalised to GAPDH from 5 independent experiments. ADAR2 levels normalised to loading control GAPDH. Statistical Analysis: Unpaired t-test; ****<0.0001. C. Representative western blots of total ADAR1 and GAPDH levels in hippocampal neurons with or without 24 h TTX treatment. D. Quantification of (C) total ADAR1 normalised to GAPDH from 8 independent experiments. Both bands were quantified. Statistical Analysis: Unpaired t-test; ns>0.05. E. Representative western blots showing total ADAR2 and GAPDH levels with increasing lengths of TTX treatment. F. Quantification of (E) total ADAR2 normalised to GAPDH from 6 independent experiments. Statistical Analysis: One Way ANOVA Dunnett’s multiple comparison test; **<0.01, ***<0.001. G. Representative western blot of nuclear ADAR2 levels in hippocampal neurons with or without 24 h TTX treatment. Cell fractionation was performed to determine the ADAR2 levels in the nucleus. Lamin B was used as a nuclear marker and RhoGDI as cytosol marker. H. Quantification of (G) nuclear ADAR2 immunoblots normalised to Lamin B from 3 independent experiments. Statistical Analysis: Unpaired T test; **<0.01. I. Representative images of hippocampal neurons with or without 24 h TTX treatment labelled with nuclear DAPI stain (blue), anti-ADAR2 (red) and anti-Fibrillarin (nucleolar marker; cyan). Bottom panels show zoom in images as indicated and the red arrows indicate cells expressing ADAR2. Scale bar=10μm. J. Quantification of (I) ADAR2 intensity per nucleus. N=3 independent dissections and n=60 cells for control and 62 cells for TTX treated. Statistical Analysis: Wilcoxon matched-pairs signed rank test, ****<0.0001. K. Analysis of percentage of cells expressing ADAR2 (I) with or without TTX. N=3 independent dissections and n=15 fields of view. Statistical Analysis: Unpaired t-test, *<0.05. L. Representative western blots of total ADAR2 and GAPDH levels in hippocampal neurons with 24 h TTX, or 24 h AP5, or both treatments, to suppress synaptic activity. M. Quantification of (L) total ADAR2 normalised to GAPDH from 4 independent experiments. ADAR2 levels normalised to loading control GAPDH. Statistical Analysis: Two-way ANOVA with Tukey’s Multiple comparisons test; ****<0.0001.

As expected, the decrease in ADAR2 following 24 h TTX treatment occurs in the nucleus (**Fig 1G,H**), where ADAR2 binds to the pre-mRNA substrates prior to mRNA splicing and maturation (Herb et al., 1996). Both the abundance of ADAR2 within cells and the proportions of cells expressing ADAR2 were decreased (**Fig 1I,J,K**). We therefore hypothesised that this activity-dependent modulation of ADAR2 could underpin the previously reported upscaling of GluK2-containing KARs in response to TTX treatment (Evans et al., 2017b).

In addition to TTX, chronic block of NMDARs with the antagonist AP5 also significantly decreased ADAR2 levels, with no additional decrease when cells were treated with both 24 h TTX and AP5 **(Fig 1L,M)**. These data demonstrate that different methods of suppressing synaptic and network activity can modulate ADAR2 levels.

### GluK2 Q/R editing in KARs is more sensitive to changes in ADAR2 levels than GluA2 Q/R editing in AMPARs

Unlike other mRNA editing sites in KAR subunits, GluK2 Q/R editing has been shown to affect KAR trafficking to the plasma membrane (Ball et al., 2010). As noted above, ADAR2 also regulates GluA2 mRNA editing and, as for GluK2, this affects the trafficking and surface expression of GluA2-containing AMPARs (Greger et al., 2003). Indeed, ADAR2 knock-out mice are deficient in both GluK2 and GluA2 editing (Higuchi et al., 2000). Moreover, GluA2 Q/R editing is modulated by changes in ADAR2 levels that occur during ischaemia (Peng et al., 2006) and in response to excitotoxic levels of glutamate (Mahajan et al., 2011). We have previously reported that 24 h TTX treatment reduces GluK2 editing and upscales KAR surface expression. We also showed that partial knockdown of ADAR2 to levels similar to those observed following 24 h TTX treatment upscaled KARs (Evans et al., 2017b). Here we investigated if mRNA editing of the AMPAR subunit GluA2 was also affected by depletion of ADAR2. To test this we used the same shRNA construct we used in our previous work that reduces ADAR2 levels to ~50% of the control and reduced the intensity of signal and percentage of cells expressing ADAR2 to levels similar to those elicited by TTX (Evans et al., 2017b). In addition, we also validated a different ADAR2 shRNA (shRNA_‘Complete’_) that ablated essentially all ADAR2 (**Fig S2**).

We then used these tools to compare how the extent of ADAR2 loss affects GluK2 and GluA2 editing. Knockdown of ADAR2 by shRNA_‘Complete’_ reduced GluK2 Q/R editing by over 60%, whereas, as reported previously (Evans et al., 2017b), shRNA_‘Partial’_ only reduced GluK2 Q/R editing by ~20%, (**Fig 2A,B**). DNA sequencing chromatographs from cells treated with shRNA_‘Complete’_ show a dramatic change in the base read of the editing site to the unedited CAG (Q) rather than edited CGG (R), while neurons treated with shRNA_‘Partial’_ show a mixture of both CGG and CAG (**Fig 2C**). Interestingly, shRNA_‘Partial’_ knockdown of ADAR2 had no effect on the Q/R editing of the AMPAR subunit GluA2 while shRNA_‘Complete’_ only reduced GluA2 editing by ~30% (**Fig 2D,E,F**). Our upscaling concomitant with a decrease in GluK2 Q/R editing (Evans et al., 2017b). Here we show additionally that, while AMPARs also undergo TTX mediated upscaling (Turrigiano, 2011), the Q/R editing status of GluA2 is not changed by TTX (Fig S3A-C) indicating that AMPAR scaling occurs via a different mechanism.

**Figure 2:**
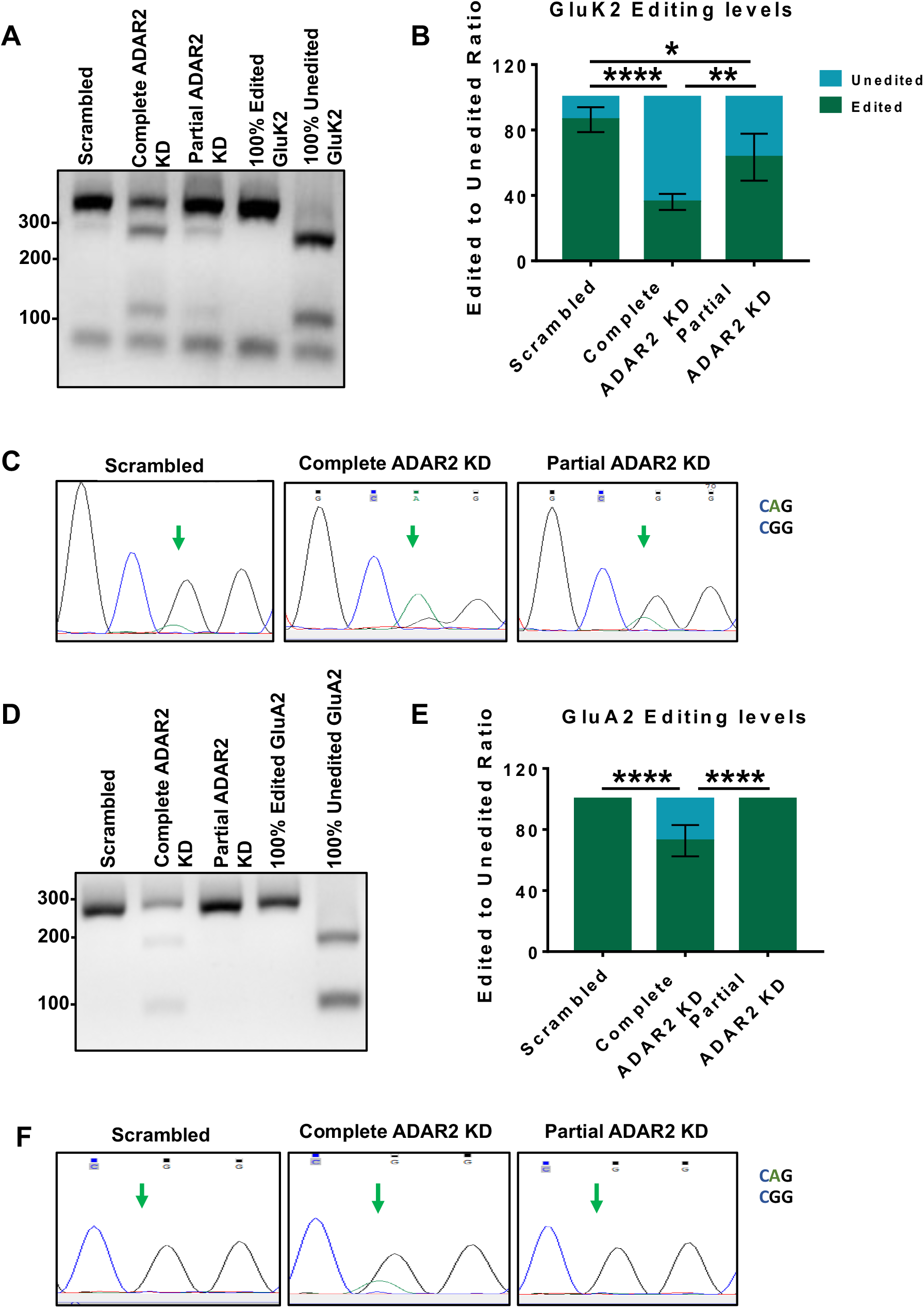
Complete and partial ADAR2 knockdown differentially alter GluK2 and GluA2 Q/R editing. A. RT-PCR and BbvI digestion analysis of GluK2 Q/R editing from hippocampal neurons infected with either scrambled, complete or partial ADAR2 KDs. B. Quantification of (A) from 4 independent experiments. Statistical Analysis: One Way ANOVA with Tukey’s multiple comparisons test; *<0.05, **<0.01, #x002A;***<0.0001. C. Sanger sequencing chromatographs of the GluK2 PCR products from hippocampal neurons infected with either scrambled, complete or partial ADAR2 KDs, showing dual A and G peaks at the editing site indicated by the green arrows. The green peak represents an A (unedited) base read and black represents a G (edited) base read. D. RT-PCR and BbvI digestion analysis of GluA2 Q/R editing from hippocampal neurons infected with either scrambled, complete or partial ADAR2 KDs. E. Quantification of (D) from 4 independent experiments. Statistical Analysis: One Way ANOVA with Tukey’s multiple comparisons test; ****<0.0001. F. Sanger sequencing chromatographs of the GluA2 PCR products from hippocampal neurons infected with either scrambled, complete or partial ADAR2 KDs. Green arrows indicate the editing site. Only samples treated with complete ADAR2 knockdown show a dual A and G peak at the editing site. The green peak represents an A (unedited) base read and black represents a G (edited) base read.

These data demonstrate that editing levels of GluK2 are selectively sensitive to changes in ADAR2 that occur as a result of TTX treatment and support a model whereby synaptic suppression-evoked loss of ADAR2 during homeostatic scaling directly promotes surface expression of GluK2-containing KARs through a reduction in GluK2 editing. We interpret this data to indicate that the Q/R editing of GluK2 subunit of KARs is more susceptible to TTX-mediated activity-dependent regulation compared to that of GluA2. This is consistent with Q/R editing of GluA2 being preferentially maintained to prevent neurotoxicity associated with Ca^2+^-permeable AMPARs and suggests that mechanisms other than ADAR2 can regulate GluA2 editing. How this is achieved is currently unclear but it is notable that ADAR1 remains unchanged during TTX treatment. Although it has been reported that ADAR1 primarily, but not exclusively, mediates R/G mRNA editing (Wong et al., 2001), we speculate that under conditions where levels of ADAR2 are diminished, ADAR1 may compensate to maintain Q/R editing of GluA2 but not GluK2. It has also been suggested that additional regulatory steps during GluA2 pre-mRNA maturation contribute to ensuring its editing levels are maintained (Penn et al., 2013).

Taken together, our data indicate that changes in levels of ADAR2 and ADAR2-mediated GluK2 Q/R editing levels provides a flexible and rapidly tunable system to control KAR forward trafficking and scaling that is not present for AMPARs.

### Partial ADAR2 knockdown mimics and occludes TTX-evoked KAR upscaling

As demonstrated in Figure 2, shRNA_‘Partial’_ reduces ADAR2 levels comparable to those following TTX treatment and results in a similar shift in the extent of GluK2 editing, while not affecting editing of the AMPAR subunit GluA2. We therefore wondered if shRNA_‘Partial’_ ADAR2 knockdown alone is sufficient to upscale KARs. Indeed, shRNA_‘Partial’_ in the absence of TTX significantly increased GluK2 surface expression with no effect on EGFR surface expression (**Fig 3A-C**). Importantly, total levels of both GluK2 and EGFR remained unaltered **(Fig S4A,B)**. Moreover, the effects of shRNA_‘Partial’_ on GluK2-KAR upscaling are reversed by rescuing levels of ADAR2 (**Fig S4C-G**). Since in our knockdown – rescue experiments there was an ‘over rescue’ of ADAR2 (**Fig S4F,G**), we tested Q/R editing of GluK2 was correspondingly increased compared to the scrambled control. Interestingly, the levels of GluK2 Q/R editing were restored to basal level (~80%) (**Fig S4E-G**) despite the over-expression of ADAR2 in the rescue condition. These results suggest that a proportion of GluK2 is resistant to Q/R editing even when excess ADAR2 is present.

**Figure 3:**
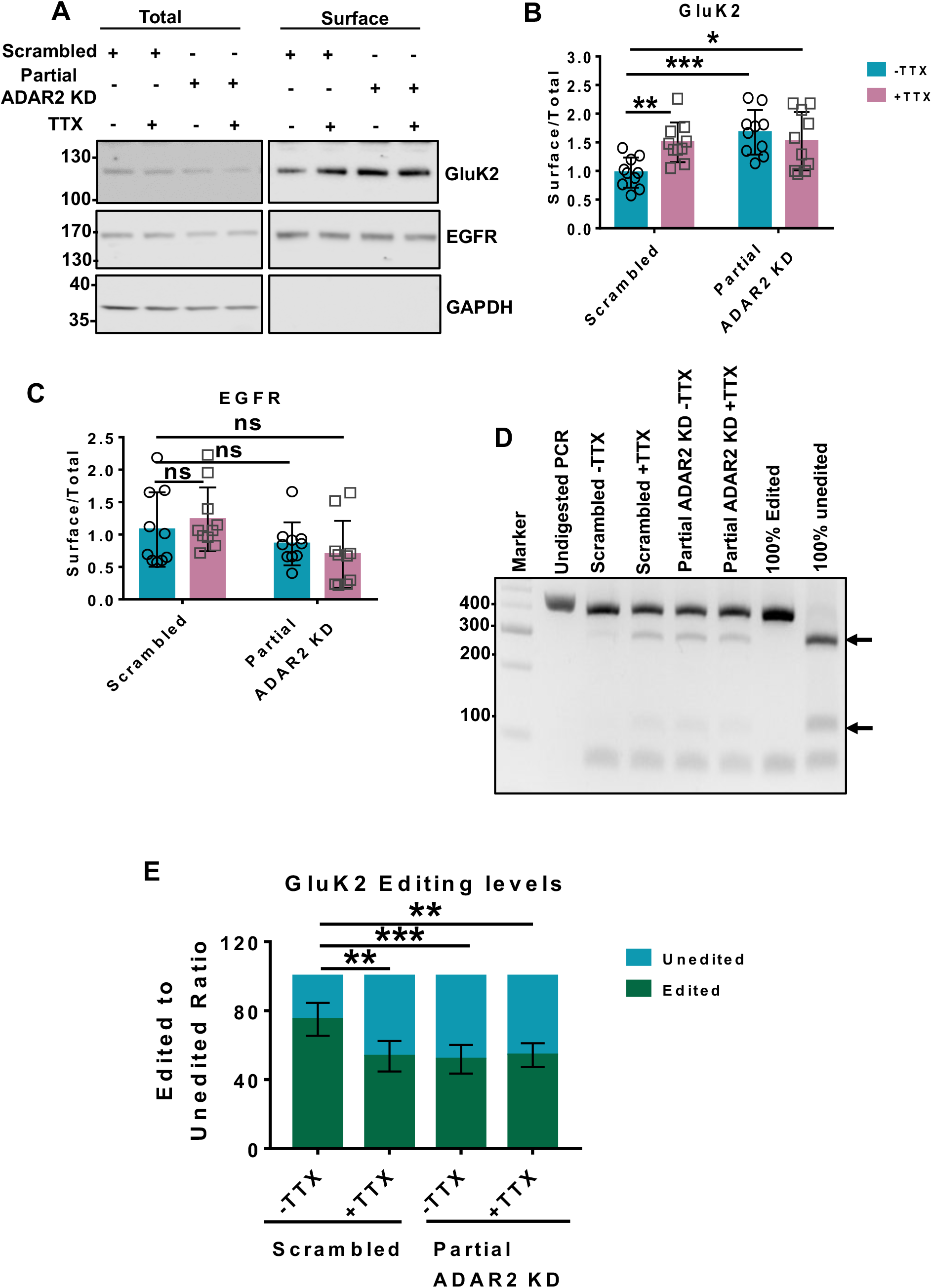
Partial ADAR2 knockdown mimics and occludes TTX-evoked KAR upscaling. A. Representative western blot of total and surface levels of GluK2, EGFR and GAPDH in scrambled or ADAR2 KD infected cells in the presence or absence of TTX. EGFR was used as a negative control while GAPDH was used as a control to show only surface proteins were labelled with biotin. B. Quantification of (A) surface levels of GluK2 from 10 independent experiments. Surface levels were normalised to their total levels. Statistical Analysis: Two-Way ANOVA with Tukey’s Multiple comparisons test; *<0.05, **<0.01, ***<0.001. C. Quantification of (A) surface levels of EGFR from 10 independent experiments. Surface levels were normalised to their total levels. Statistical Analysis: Two-Way ANOVA with Tukey’s Multiple comparisons test; ns>0.05. D. RT-PCR and BbvI digestion analysis of GluK2 Q/R editing from hippocampal neurons infected with either scrambled or partial ADAR2 KD with or without TTX. E. Quantification of (D) from 5 independent experiments. Statistical Analysis: One Way ANOVA with Tukey’s multiple comparisons test; **<0.01, ***0.001.

Surprisingly, complete ablation of ADAR2 with shRNA_‘Complete’_ had no effect on GluK2 surface expression (**Fig S5A,B**), suggesting the effect of physiologically relevant partial loss of ADAR2 differ from that of the complete knockdown. It is possible that upon complete loss of ADAR2 compensatory mechanisms exist to restore cellular homeostasis, and it is also important to note that complete ablation of ADAR2 leads to further reduction in GluK2, as well as GluA2, editing, making these results difficult to interpret.

Importantly, application of both TTX and shRNA_‘Partial’_ was not additive (**Fig 3A,B**) and did not further decrease GluK2 Q/R editing compared to each individual treatment alone (**Fig 3D,E**). The fact that partial ADAR2 knockdown is sufficient to upscale KARs and that the effects of TTX are occluded by shRNA_‘Partial’_ provide further support for the proposal that TTX-induced GluK2 upscaling is mediated by a reduction in ADAR2 levels.

### TTX promotes proteasomal degradation of ADAR2

We next explored the mechanisms underlying ADAR2 loss during scaling. As shown in **Fig 4A**, TTX does not alter ADAR2 mRNA levels indicating that transcriptional changes are not involved so we investigated possible mechanisms for activity-dependent ADAR2 degradation.

**Figure 4:**
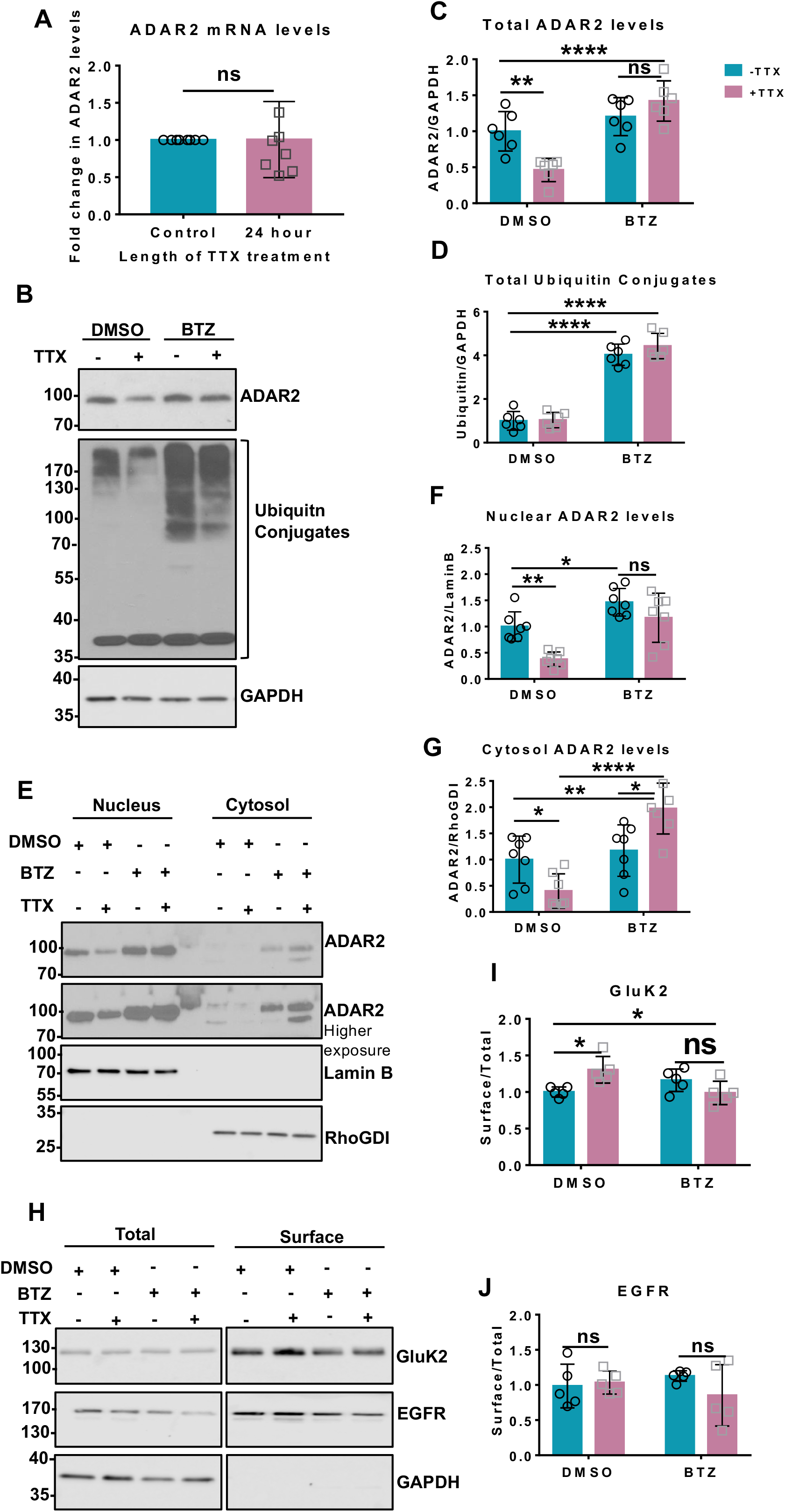
TTX promotes proteasomal degradation of ADAR2. A. RT-qPCR analysis of mRNA levels of ADAR2 post TTX treatment showing no changes in the ADAR2 mature mRNA transcripts from 7 independent experiments. Statistical analysis: Unpaired t-test; ns>0.05. B. Representative western blots of total ADAR2, total ubiquitin and GAPDH levels in neurons treated with either DMSO or 1μM Bortezomib (BTZ) for 20 h either in the presence or absence of 24 h TTX. C. Quantification of total ADAR2 immunoblots normalised to GAPDH from 6 independent experiments. Statistical Analysis: Two-way ANOVA with Tukey’s multiple comparisons test: **<0.01, ****<0.0001, ns>0.05. D. Quantification of total ubiquitin conjugated products normalised to GAPDH from 6 independent experiments. Statistical Analysis: Two-way ANOVA with Tukey’s multiple comparisons test: ****<0.0001. E. Representative western blots of ADAR2 in the nucleus and cytoplasm in the presence of DMSO or BTZ with or without TTX treatment. Lamin B was used as a nuclear marker and RhoGDI as cytosol marker. F. Quantification of nuclear ADAR2 from 7 independent experiments. Nuclear ADAR2 levels were normalised to Lamin B. Statistical Analysis: Two-way ANOVA Tukey’s multiple comparisons test: *<0.05, **<0.01, ns>0.05. G. Quantification of cytosolic ADAR2 from 7 independent experiments. Cytosolic ADAR2 was normalised to RhoGDI level. Statistical Analysis: Two-way ANOVA with Tukey’s multiple comparisons test: *<0.05, **<0.01, ****<0.0001. H. Representative western blots of total and surface levels of GluK2, EGFR and GAPDH in DMSO and BTZ (20 h, 1μM) treated cells in the presence or absence of 24 h TTX. EGFR was used as a negative control while GAPDH was used as a control to show only surface proteins were labelled with biotin. I. Quantification of surface levels of GluK2 from 5 independent experiments. Surface levels were normalised to their total levels. Statistical Analysis: Two-Way ANOVA with Tukey’s Multiple comparisons test; *<0.05, ns>0.05. J. Quantification of surface levels of EGFR from 5 independent experiments. Surface levels were normalised to their total levels. Statistical Analysis: Two-Way ANOVA with Tukey’s Multiple comparisons test; ns>0.05.

The nuclear protein Pin1 retains ADAR2 in the nucleus to prevent its export to the cytosol where it is ubiquitinated and degraded (Marcucci et al., 2011). It has also been reported that Pin1-mediated stabilisation is an important regulator of ADAR2 editing activity during development in cortical neurons (Behm et al., 2017). We therefore wondered if destabilisation of the Pin1-ADAR2 interaction underpins the TTX-mediated ADAR2 loss. However, Pin1 levels were unchanged following TTX treatment (**Fig S6A,B**).

ADAR2 phosphorylation at threonine 32 (T32) has also been reported to be crucial for the ADAR2-Pin1 interaction (Marcucci et al., 2011) so we made phosphonull (T32A) and phosphomimetic (T32D) ADAR2 mutants. ADAR2(T32D) binds very strongly to Pin1 in GFP-trap assays compared to WT and ADAR2(T32A) (**Fig S6C,D**). We therefore tested if the phosphonull or phosphomimetic ADAR2 mutants were more sensitive to TTX treatment. We first knocked down endogenous ADAR2 and replaced it with HA-tagged WT ADAR2 (**Fig S6E,F**). More than 80% of the cells expressed this ADAR2 knockdown-rescue protein, similar to the percentage of scrambled treated neurons that express endogenous ADAR2. We then investigated the stability of the phosphonull or phosphomimetic ADAR2 mutants in response to TTX treatment. Similar to WT ADAR2, levels of both mutants were significantly decreased by TTX treatment (**Fig S6G,H**). Since both phosphonull and phosphomimetic mutants of ADAR2, which decrease or enhance binding to Pin1 respectively, were equally susceptible to the TTX-mediated loss these experiments suggest that alterations in the Pin1-ADAR2 interaction do not underpin ADAR2 loss during TTX mediated upscaling.

We next determined the effects of TTX on ADAR2 stability in the presence or absence of the proteasomal inhibitor Bortezomib (BTZ) (Chen et al., 2011). BTZ prevented the TTX-evoked decrease in ADAR2 (**Fig 4B,C**) and resulted in the accumulation of ubiquitinated proteins (**Fig 4B,D**). We performed nuclear and cytoplasmic fractionation experiments to determine if ADAR2 is exported from the nucleus for degradation in the cytosol. BTZ prevented the TTX-evoked decrease in ADAR2 in both the nuclear and cytosolic fractions, and actually led to a significant accumulation of ADAR2 in the cytosol (**Fig 4E,F,G**). Thus, ADAR2 may be exported to the cytosol for ubiquitination, ubiquitinated in the nucleus and exported to the cytosol, or a combination of both. While the exact mechanisms remain to be determined, these experiments show that suppression of synaptic activity induces ADAR2 ubiquitination and proteasomal degradation.

Since BTZ prevents the loss of ADAR2 by TTX treatment, we next tested whether BTZ also blocks KAR upscaling. Indeed, surface biotinylation showed that BTZ prevents TTX-induced increases in surface expressed GluK2 (**Fig 4H,I**) with no effect on EGFR (**Fig 4H,J**). These results support the hypothesis that proteasomal degradation of ADAR2 following TTX treatment is both necessary and sufficient for KAR upscaling.

Taken together our data demonstrate that suppression of synaptic activity reduces ADAR2 leading to decreased KAR editing which, in turn, directly mediates KAR upscaling via increased KAR assembly and ER exit of unedited GluK2(Q) compared to edited GluK2(R) (**Fig 5**). These results show that regulation of ADAR2 stability and changes in GluK2 editing underpin a novel and specific mechanism to tune the surface expression of KARs. Given that KARs play many roles in controlling neuronal network activity (Contractor et al., 2011; Evans et al., 2017a), that they have recently been identified as inducers of AMPAR plasticity (Petrovic et al., 2017), and that their dysfunction has been implicated in a number of neurological disorders (Crepel and Mulle, 2015; Lerma and Marques, 2013), it is likely that ADAR2 mediated control of KAR surface expression plays a wide role in neuronal function and dysfunction.

**Figure 5:**
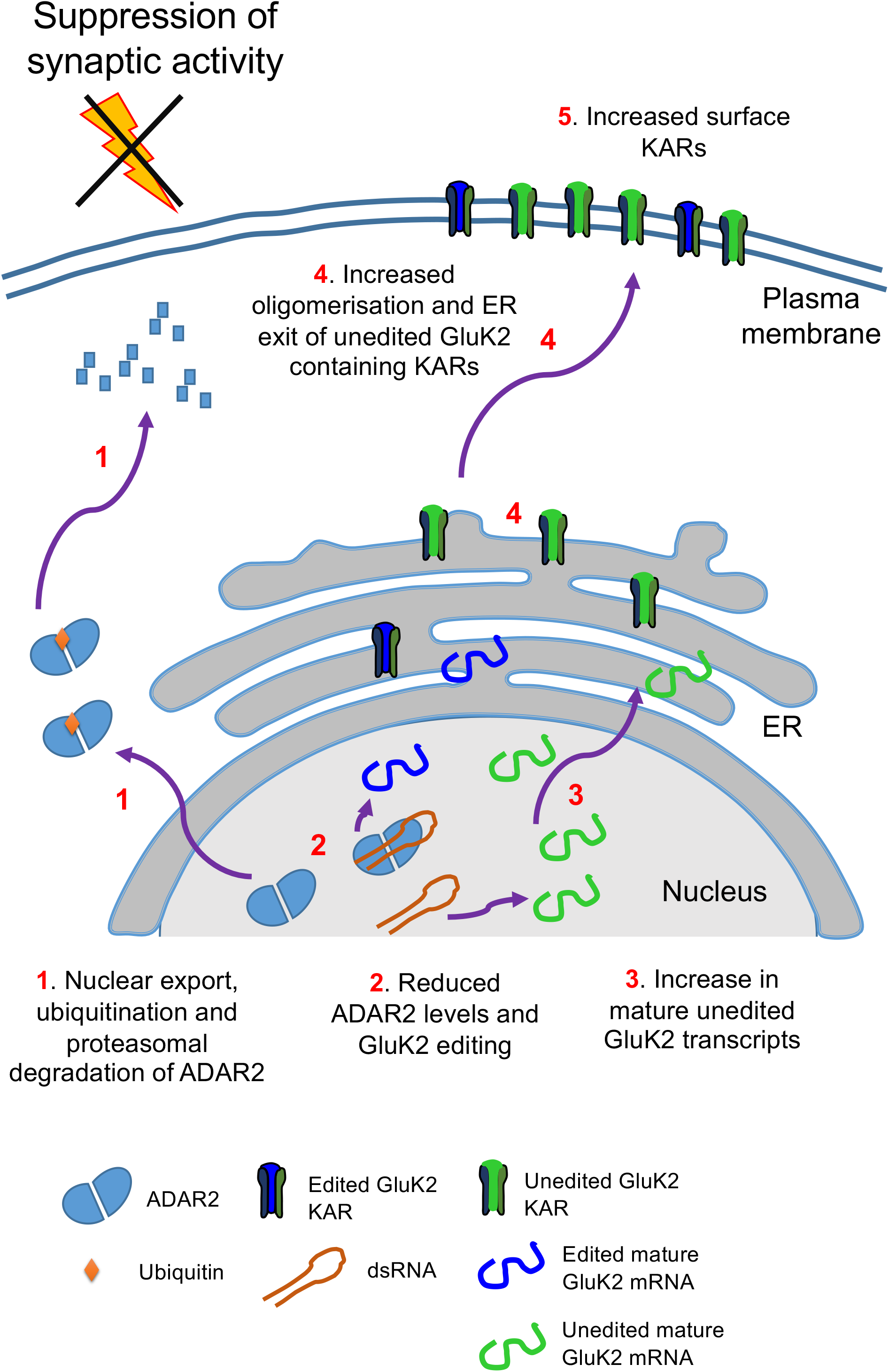
Schematic of ADAR2 mediated Q/R editing regulating GluK2 containing KARs homeostatic upscaling. Under basal conditions unedited GluK2 transcripts are edited at their Q/R site by ADAR2 resulting in ~80% of mature GluK2 transcripts being edited. The resultant edited and unedited GluK2 subunits oligomerise in the ER and traffic to the surface. Under conditions of synaptic activity suppression with TTX treatment, ADAR2 undergoes proteasomal degradation in the cytosol (1). This results in less ADAR2 editing of GluK2 pre-mRNA transcripts (2) and increased levels of unedited mature GluK2 transcripts (3). The subsequent increase in the proportion of unedited GluK2(Q) allows enhanced oligomerisation and ER exit (4) to increase surface expression of GluK2-containing KARs on the surface (5).

## Materials and Methods

### Primary neuronal cultures

Primary rat hippocampal neurons were dissected from E18 Wistar rat pups as previously described (Rocca et al., 2017). Briefly neurons were dissected from E18 Wistar rats followed by trypsin dissociation and cultured for up to 2 weeks. For the first 24 h, cells were grown in plating media: Neurobasal media (Gibco) supplemented with 10% horse serum (Sigma), B27 (1x, Gibco), P/S (100 units penicillin and 0.1mg/ml streptomycin; ThermoScientific) and 5mM Glutamax (Gibco). After 24 h, plating media was replaced with feeding media (Neurobasal containing 2mM Glutamax and lacking horse serum and P/S). For biochemistry experiments, cells were plated at a density of 500,000 per 35mm well and 250,000 per coverslip for imaging experiments.

#### ADAR2 cloning

ADAR2 was cloned from rat neuronal cDNA and ADAR2 shRNA knockdown and knockdown-rescue viruses were generated as previously described (Rocca et al., 2017). ADAR2 was cloned from rat neuronal cDNA into the KpnI and XbaI sites of the vector pcDNA3 with a HA tag at its N-terminus. Phosphomutants of ADAR2 were generated by site-directed mutagenesis. Pin1 was cloned from rat neuronal cDNA into the EcoRI and BamHI sites of the vector pEGFP-N1.

#### Lentivirus generation

For ADAR2 knockdown experiments, shRNA sequences targeting ADAR2 cloned into a modified pXLG3-GFP vector (Rocca et al., 2017) under the control of a H1 promoter. The ADAR2 target sequences were:

~~~
Complete shRNA     AAGAACGCCCTGATGCAGCTG
Partial shRNA         AACAAGAAGCTTGCCAAGGCC
~~~

For rescue experiments, shRNA-insensitive HA-ADAR2 was cloned into a modified pXLG3-GFP vector under the control of an SFFV promoter.

The viruses were produced in HEK293T cells as reported previously (Rocca et al., 2017), harvested and added to DIV 9/10 hippocampal neurons for 5 days and lysed accordingly. For 24 h treatment experiments, cells were treated with on the 4^th^ day after virus addition and harvested accordingly on the 5^th^ day after the completion of the time course.

#### Scaling, developmental and TTX timecourse and BTZ treatment

For scaling experiments, cells were treated with 1μM TTX (Tocris) for 24 h and were either lysed directly in 1x sample buffer and heated for 10 min at 95°C or were used for either surface biotinylation or fractionation experiments (see below). For the TTX timecourse, cells were harvested directly into 1x sample buffer (4 x sample buffer (0.24M Tris-HCL, 8% SDS, 40% glycerol, 10% β-mercaptoethanol and 0.009% Bromophenol blue) diluted in water). In experiments inhibiting proteosomal degradation, cells were treated with bortezomib (BTZ; Cell Signalling) dissolved in DMSO for 20 h at 1μM concentration. The control cells were treated with an equal volume of DMSO.

#### Cell surface biotinylation and streptavidin pulldown

Cell surface biotinylation was performed essentially as previously described (Evans et al., 2017b). All steps were performed on ice with ice-cold buffers unless stated otherwise. Live hippocampal neurons post stated treatments were washed twice in phosphate buffered saline (PBS). Surface proteins were labelled with membrane impermeable Sulfo-NHS-SS biotin (0.3mg/ml, Thermo Scientific) for 10 min on ice and washed 3x with PBS. 100mM NH_4_Cl was added to quench free biotin-reactive groups and cells were extracted with lysis buffer (50mM Tris pH 7.4, 150mM NaCl, 1% triton, 0.1% SDS, protease inhibitors (Roche)), incubated on ice for 30 min and centrifuged (15,000g, 4°C, 20 min) to remove cell debris. For streptavidin pulldown, each lysate was added to 30μl of streptavidin beads (Sigma) and left on a wheel to rotate for 90 min at 4°C. The beads were then washed 3x with wash buffer (lysis buffer without protease inhibitors) and proteins eluted with 2x sample buffer and boiled for 10 min at 95°C. The samples were then resolved by SDS-PAGE and immunoblotted.

#### Subcellular fractionation

All the steps were performed on ice with ice-cold buffers unless stated otherwise. Following stated treatments, the cells were washed with PBS followed by addition of buffer 1 (150mM NaCl, 50mM HEPES pH 7.4, 25μg/ml Digitonin and protease inhibitors), incubated for 20 min, scraped, homogenised and centrifuged at 15000g for 30 min at 4°C. The supernatant consisted of the cytosolic proteins, while the pellet was resuspended in buffer 2 (Buffer 1 with 1% triton), incubated for 20 min and again centrifuged at 15000g for 30 min at 4°C. The supernatant consisted of mitochondrial proteins while the pellet was resuspended in buffer 3 (150mM NaCl, 50mM HEPES pH 7.4, 0.5% sodium deoxycholate, 0.1% SDS, protease inhibitors and 0.5% triton), incubated for 1 h and centrifuged at 15000g for 30 min at 4°C. The supernatant was discarded and the pellet, consisting of nuclear proteins, was resuspended in buffer 3. The cytosolic supernatant was concentrated using 4 volumes of acetone (kept at −20°C), incubated at −20°C for 1 h and spun for 20 min at 1500g and resuspended in buffer 3. BCA assay was then performed to determine protein concentrations and allow equal loading.

#### BCA Assay

BCA Assay was performed using a commercial kit (Pierce, Thermo Scientific) following manufacturer’s instructions. Following 30 min incubation at 37°C, the samples were read using a plate reader (Versamax Microplate reader, Molecular devices) at a wavelength of 562nm.

#### Western blotting and antibodies

Antibodies used: GluK2 (1:1000, Millipore, rabbit polyclonal), GluA2 (1:1000, Synaptic Systems, rabbit polyclonal), ADAR2 (1:1000, Sigma, rabbit polyclonal), GAPDH (1:10,000, Abcam, mouse monoclonal), RhoGDI (1:1000, Abcam, rabbit polyclonal), Lamin B (1:1000, Santa Cruz, Goat polyclonal), EGFR (1:1000, Abcam, rabbit polyclonal), HA (1:2000, mouse, Sigma), Pin1 (mouse monoclonal, 1:1000, Santa Cruz), ADAR1 (mouse monoclonal, 1;1000, Santa Cruz) and GFP (rat monoclonal, 1:10,000, Chromotek). Western blots were imaged and quantified using LI-COR Image Studio software or developed on X-ray film in a dark room using developer and fixer solutions. The blots were then scanned and quantified using FIJI ImageJ studio. Surface levels were normalised to their respective total levels. Nuclear protein levels were normalised to LaminB and cytosolic protein levels to RhoGDI. Treated samples were normalised to their control samples.

#### RNA extraction, RT-PCR, and BbvI digestion and RT-qPCR

RNA samples were extracted from DIV14/15 hippocampal neurons following the stated treatments using RNeasy Mini Kit (Qiagen) according to the manufacturer’s instructions. 1μg of RNA was used per condition and reverse transcribed to cDNA using RevertAid First Strand cDNA Synthesis Kit (Thermo Scientific) following the manufacturer’s instructions. The following primers (spanning the M2 region of GluK2 (Bernard et al., 1999) and GluA2) were used, giving a PCR product of 452bp and 252bp:

GluK2 F: 5’-GGTATAACCCACACCCTTGCAACC-3’
GluK2 R: 5’-TGACTCCATTAAGAAAGCATAATCCGA-3’
GluA2 F: 5’-GTGTTTGCCTACATTGGGGTC-3’
GluA2 R: 5’-TCCTCCTACACGGCTAACTTA-3’

5μl of cDNA was used to set up PCR reactions [50μl total, 35 cycles, 20s denaturing at 95°C, 10s annealing at 60°C and 15 s elongation at 70°C].

To determine the level of RNA editing, BbvI (New England Biolabs) digestion was used as previously (Bernard et al., 1999). Total 20μl digestion was set up using 10μl of PCR product at 37°C for 2 h. All of the digested product was run on 4% agarose gel and the ethidium bromide stained bands were imaged using UV transilluminator and quantified using FIJI NIH ImageJ. To determine the level of editing in GluK2, the following formula was used: [Intensity of 376 (edited)/Intensity of (376 (edited) + 269 (unedited))]*100. The band at 76bp allowed to determine equal loading. For GluA2, [Intensity of 158 (edited)/Intensity of (158 (edited) + 94 (unedited))]*100. Purified PCR products were also sent for sequencing to Eurofins Genomics at 4ng/μl along with the above GluK2 and GluA2 F primers, to obtain sequence chromatographs.

For RT-qPCR, 2μl of the cDNA samples per condition were mixed with PowerUp SYBR green Master Mix (Life Technologies) and forward and reverse primers targeting ADAR2 and GAPDH and amplified quantitatively using Real Time PCR System (MiniOpticon, BioRAD) for 40 cycles and Ct values were recorded. Each reaction was performed in triplicate and average Ct was measured per condition. ADAR2 Ct values were normalised to GAPDH Ct values and ADAR2 mRNA fold difference value of TTX treated conditions was normalised against the untreated control. Melting curve of the primers were also determined to ensure the specificity of the primers and lack of primer dimer formation. The primers used were: ADAR2 F – TCCCGCCTGTGTAAGCAC, GAPDH AATCCCATCACCATCTTCCA.

#### GFP trap

GFP-trap protocols were as previously published (Guo et al., 2017). HEK293T cells were transfected the next day using Lipofectamine^TM^3000 and 2.5μg of each construct. 48 h post-transfection, cells were washed with PBS and lysed and harvested in lysis buffer (20mM Tris pH7.4, 137mM NaCl, 2mM sodium pyrophosphate, 2mM EDTA, 1% triton X-100, 0.1% SDS, 25mM β-glycerophosphate, 10% glycerol, protease inhibitors (Roche), phosphatase inhibitor cocktail 2 (1:100, Sigma)). The lysates were left to incubate for 30 min on ice and centrifuged at 1500g for 20 min at 4°C to remove any cell debris. The supernatant was then added to 5μl of GFP-trap beads (Chromotek), incubated on wheel at 4°C for 90 min and washed 3x with wash buffer (lysis buffer without protease or phosphatase inhibitors). The samples were then lysed in 2x sample buffer, heated at 95°C for 10 min and separated using SDS-PAGE.

#### Fixed immunostaining, imaging and analysis

Immunostaining was performed as previously described (Glebov et al., 2015). For fixed immunostaining, cells post TTX treatment or lentiviral treatment as indicated were fixed with 4% formaldehyde for 10 min, washed 3 times with PBS, treated with 100mM glycine to quench any remaining formaldehyde and washed 3 times with PBS. The cells were permeabilised and blocked with 3% BSA in PBS and 0.1% triton for 30 min. The cells were then incubated for 1 h with primary antibodies (anti-ADAR2 (Abcam, rabbit) 1:400, anti-Fibrillarin (Abcam, mouse) 1:400 and anti-HA (Sigma, mouse) 1:600) in 3% BSA at room temperature, washed 3 times for 5 min each with PBS and incubated for 45 min with the indicated secondary antibodies (Jackson Immunoresearch Antibodies, 1:400) in 3% BSA at room temperature. Three x 5-minute washes were performed with PBS and the cells were mounted using DAPI containing fluoromount.

A Leica SP5-II confocal laser scanning microscope attached to a Leica DMI 6000 inverted epifluorescence microscope was used to image the coverslips. The confocal images were captured under 63x objective, with 1024×1024 pixel resolution and 1xoptical zoom. Frame average of 2 was taken with a Z-stack of 6-8 Z-planes with 0.5μM interval.

FIJI NIH Image J was used to compress the z-stacks and analyse the mean intensity per nucleus using the DAPI channel to draw regions of interest. To calculate the percentage of cells expressing ADAR2, all the cells expressing ADAR2 were manually counted per image taken.

#### Statistical Analysis

Mean value were calculated for all data and all error bars show standard deviation. All statistical analysis was performed using GraphPad Prism software version 7.0 as stated. N= Number of dissections and n= number of cells. Unpaired t-test was performed when comparing changes between two different groups. One-way Analysis of Variance (ANOVA) was performed to compare mean changes within more than two groups and Two-way ANOVA was used to compare mean differences between multiple groups with two independent variables. Dunnett’s multiple post-test was performed to determine any significant changes when compared to the control group while Tukey’s multiple post comparisons were performed to compare multiple groups at a time.

## Acknowledgements

We are grateful to the MRC, BBSRC and Wellcome Trust for financial support. SG and AJE were supported by Wellcome Trust Dynamic Cell Biology PhD studentships. We thank Dr Yasuko Nakamura for excellent technical and logistical support.

## Author Contributions

SG performed all of the experiments. AJE and KAW provided specialised constructs, advice and assisted in some experiments. JMH supervised the research. All authors contributed to writing the manuscript.

---

**Figure S1: TTX does not alter total levels of KAR subunits.**

A. Representative western blots of total GluK2, GluK5 and GAPDH levels in hippocampal neurons treated with or without 24 h TTX.

B. Quantification of (D) total GluK2 normalised to GAPDH from 5 independent experiments. Statistical Analysis: Unpaired t-test; ns>0.05.

C. Quantification of (D) total GluK5 normalised to GAPDH from 5 independent experiments. Statistical Analysis: Unpaired t-test; ns>0.05.

**Figure S2: Validation of complete and partial ADAR2 knockdown.**

A. Representative western blots of total ADAR2 and GAPDH levels in hippocampal neurons treated with either scrambled, complete or partial ADAR2 KDs.

B. Quantification of total ADAR2 normalised to GAPDH from 5 independent experiments for scrambled and partial knockdown and 4 independent experiments for complete knockdown. Statistical Analysis: One-Way ANOVA with Tukey’s multiple comparisons test; *<0.05, ***<0.001, ****<0.0001.

C. Representative images of hippocampal neurons imaged for DAPI (blue), GFP (green, lentivirus infected cells), ADAR2 (red) and Fibrillarin (nucleolar marker; cyan) for cells infected with either scrambled, complete or partial ADAR2 KD. Bottom panels show zoom in images as indicated and the red arrows indicate cells expressing ADAR2. Scale bar=10μm.

D. Quantification of ADAR2 intensity per nucleus. N=3 independent dissections and n=75 cells (complete KD), 86 cells (scrambled) and 99 cells (Partial knockdown). Statistical Analysis: One Way ANOVA with Tukey’s multiple comparisons test, ***<0.001, ****<0.0001.

E. Quantification of percentage of cells expressing ADAR2. N=3 independent dissections and n=11-13 fields of view. Statistical Analysis: One Way Anova with Tukey’s multiple comparisons test, ****<0.0001.

**Figure S3: TTX does not alter GluA2 editing.**

A. Schematic of BbvI digestion analysis on the PCR amplified M2 region of GluA2.

B. RT-PCR and BbvI digestion analysis of GluA2 Q/R editing from hippocampal neurons treated with or without TTX.

C. Sanger sequencing chromatographs of the GluA2 PCR products from hippocampal neurons treated with or without TTX, showing no changes in editing levels. Black arrows indicate the editing site. Green peak represents an A (unedited) base read and black represents a G (edited) base read.

**Figure S4: Partial loss of ADAR2 does not change total levels of GluK2 and effect of the partial loss can be rescued upon ADAR2 addition.**

A. GluK2 total levels remain unaltered following partial ADAR2 KD and TTX treatment from 9 independent experiments. Statistical Analysis: Unpaired t-test; ns>0.05.

B. EGFR total levels remain unaltered following partial ADAR2 KD and TTX treatment from 5 independent experiments. Statistical Analysis: Unpaired t-test; ns>0.05.

C. Representative western blot of total and surface levels of GluK2, EGFR and GAPDH in scrambled or Partial ADAR2 KD or Partial ADAR2 KD with WT ADAR2 rescue infected cells. EGFR was used as a negative control while GAPDH was used as a control to show only surface proteins were labelled with biotin.

D. Quantification of (C) surface levels of GluK2 from 5 independent experiments. Surface levels were normalised to their total levels. Statistical Analysis: One-Way ANOVA with Tukey’s Multiple comparisons test; ***<0.001, ****<0.0001.

E. Quantification of (C) surface levels of EGFR from 5 independent experiments. Surface levels were normalised to their total levels. Statistical Analysis: One-Way ANOVA with Tukey’s Multiple comparisons test; ns>0.05.

F. Representative western blot of total levels of ADAR2 and GAPDH in scrambled or Partial ADAR2 KD or Partial ADAR2 KD with WT ADAR2 rescue infected cells.

G. Quantification of (F) total levels of ADAR2 normalised to GAPDH from 5 independent experiments. Statistical Analysis: One-Way ANOVA with Tukey’s Multiple comparisons test; *<0.05, ***<0.001.

H. Representative images of the PCR products separated on a 4% agarose gel post BbvI digestion to determine edited to unedited GluK2 ratio. 100% edited and unedited GluK2 constructs were used as a control to ensure BbvI cut activity and validity of the assay.

I. Quantification of percentage of edited to unedited ratio of GluK2 population present as shown in A. n=4 independent dissections; One-way ANOVA with Tukey’s multiple comparisons test; *p<0.05, **p<0.01, ****p<0.0001.

J. Representative chromatographs of PCR products comparing scrambled infected cells with the partial knockdowns and partial knockdown with WT ADAR2 rescue infected cells at the Q/R editing site of GluK2. The undigested PCR products were sent for sequencing to determine changes in the dual peaks obtained at the site of editing as indicated by the green arrow. A peak (green) represents the unedited base while the G peak (black) represents the edited base. The image is representative of 3 repeats.

**Figure S5: Complete loss of ADAR2 results in no overall changes in GluK2 surface expression.**

A. Representative western blots of surface and total GluK2 and GAPDH levels in hippocampal neurons treated with either scrambled, complete or partial ADAR2 KDs.

B. Quantification of surface levels of GluK2 from 5 independent experiments. Surface levels were normalised to their total levels. Statistical Analysis: One-Way ANOVA with Tukey’s Multiple comparisons test; ns>0.05, ***<0.001, ****<0.0001. ns>0.05.

**Figure S6: TTX-mediated scaling is not dependent on Pin1 or ADAR2 phosphorylation.**

A. Representative western blots of total Pin1 and GAPDH levels in neurons with or without TTX treatment for 24h.

B. Quantification of (A) Pin1 levels normalised to GAPDH from 9 independent experiments. Statistical Analysis; Unpaired t-test: ns>0.05.

C. Representative blots of GFP-trap performed in HEK293T cells where Pin-GFP was overexpressed with either HA-WT ADAR2, HA-T32D ADAR2 or HA-T32A ADAR2. The pulldowns were blotted for HA, GFP and GAPDH. Free GFP was overexpressed with WT-ADAR2 as a negative control.

D. Quantification of (C) showing interaction of HA-T32D ADAR2 with Pin1-GFP. The HA signal was normalised to the GFP control. N=4 independent experiments. Statistical Analysis: One-way ANOVA with Tukey’s multiple comparisons test; ***<0.001, ****<0.0001.

E. Representative confocal images showing neurons imaged for DAPI (nucleus), GFP to represent the cells infected with ADAR2 knockdown, anti-ADAR2 to show successful rescue of ADAR2 in the same cells infected with the knockdown and anti-HA to show the rescues were HA tagged.

F. Quantification showing the percentage of the cells expressing the ADAR2 rescue is comparable to the number of control (scrambled) cells expressing endogenous ADAR2 (E). N=4 independent dissections and n=12-14 fields of view. Statistical Analysis: Unpaired t-test; ns>0.05.

G. Representative western blots of total ADAR2 and GAPDH in cells infected with either scrambled or complete KD or complete KD rescue lentiviruses expressing either WT ADAR2, T32D ADAR2 or T32A ADAR2, in the presence or absence of TTX.

H. Quantification of (G) total ADAR2 levels normalised to GAPDH from 7 independent experiments. Each TTX treated condition was normalised to their respective non-treated control. Statistical Analysis: Unpaired t-test; *<0.05, **<0.01, ***<0.00, ****<0.0001.

